# Conserved Residues in the Gα interface show subtype specificity in Gβγ coupling

**DOI:** 10.64898/2026.01.29.702674

**Authors:** Wenyuan Wei, H. Dalton Taylor, Ning Ma, Andrei S. Rodin, Sergio Branciamore, Henrik G. Dohlman, Nagarajan Vaidehi

## Abstract

Although the structural basis of selective G-protein coupling to G protein coupled receptors (GPCRs) is well characterized, the mechanisms underlying selective interactions between distinct Gα subtypes (Gαs, Gαi, Gαq, and Gα12/13) and Gβγ remain poorly understood. While conserved residues in Gα subtypes are often assumed to have similar functions, they may instead modulate coupling selectivity by altering the frequency and stability of contacts at the Gα:Gβγ interface. Using molecular dynamics (MD) simulations combined with the interpretable machine learning method, Bayesian Network Model (BNM), and protein-protein proximity (BRET) assays, we show that conserved residues in the two closely related Gαi/o and Gαq/11 subfamilies contribute differentially to Gβγ coupling. These conserved residue “hotspots” on Gαi and Gαq produced divergent functional effects on Gβγ coupling, indicating that conservation does not ensure functional equivalence. These findings suggest that local microenvironment and paralog-specific allosteric coupling shape how conserved interface residues contribute to protein-protein coupling. The framework provides a systematic approach for dissecting subtype-specific mechanisms, with implications for drug design and for annotating the functional relevance of disease-associated variants. The computational methods used here are broadly applicable to other homologous protein families.

## Introduction

Analyses of protein paralogs provide a direct test of whether conserved residues play a similar role in functionally similar proteins. This is a central tenet of the sequence–structure– function paradigm. Although the assumption that conserved residues contribute similarly to protein-protein interactions holds in some cases, reported exceptions indicate that conservation alone does not guarantee functional equivalence^1,2^. These observations suggest that local microenvironments and long-range allosteric communication can modulate how individual residue positions contribute to signaling outputs. Previous studies have shown that conserved residues in the GPCR:Gα interface can indeed contribute differentially to Gα subtype coupling by GPCRs^3^. Here we consider conserved residues that contribute to coupling of Gα subtypes to Gβγ. The Gα subunit of the trimeric G protein is a GTPase that dissociates from Gβγ upon activation and engages downstream pathways. Humans encode 16 Gα subtypes, functionally categorized into four subfamilies (Gαs, Gαi/o, Gαq/11, Gα12/13), and these couple with differential strengths to receptors. However, it is not known whether conserved residues in Gα subtypes contribute to coupling with differential strength to Gβγ.

Structurally, Gα proteins consist of two domains: a Ras-like domain with GTPase activity and an α-helical domain (AHD) that participates in nucleotide binding. Within the Ras-like domain, three switch regions (SWI–III) rearrange upon GTP binding, promoting Gα-Gβγ dissociation^1^. The nucleotide binding pocket lies between the Ras domain and the AHD, which undergoes opening upon GPCR mediated activation of Gα. Despite broad conservation in sequence and structure, Gα subfamilies trigger distinct effector pathways: Gαs activates adenylate cyclase to elevate cAMP^4^; Gαq/11 stimulates phospholipase Cβ isoforms to produce intracellular Ca^2^□ signals^5^; Gα12/13 activates RhoGEFs to regulate Rho-family GTPases^6^; and Gαi/o inhibits adenylate cyclase and activates GIRK channels^4,7^.

Due to the larger number of distinct GPCRs in comparison to the number of G protein subtypes and downstream effectors, G proteins represent a critical bottleneck in GPCR-mediated signaling pathways. Perturbations of G protein function, especially that of Gα, can therefore impact multiple signaling pathways. Indeed, many reported Gα mutations are deemed disease associated. For example, Gαs substitutions at Arg201 (e.g., R201C), which slow GTPase activity, are linked to McCune–Albright syndrome and various cancers^8,9^. More than 200 unique variants of Gαo are linked to epileptic encephalopathy; these mutations are scattered across the AHD and Ras-like domain, and fall into multiple mechanistically distinct groups ^10^. In our previous study^1^ we showed that one of these mutations, in a conserved arginine in an evolutionarily conserved Gly-Arg-Glu triad (“G-R-E motif”), impede Gβγ subunit dissociation. We showed that this triad forms an allosteric link between the γ-PO_4_ group of GTP and the Gβγ interface on Gα. Substitutions at a conserved “catalytic glutamine” essential for GTP hydrolysis, in Gαq or Gα11 are found in nearly all cases of uveal melanoma ^11^. Hewitt-Valentin et al showed that different substitutions at the catalytic Gln produce distinct active state conformations of Gα^11^. That study showed that Gα function arises from an ensemble of active states, some of which are preferentially selected in disease and therefore may be differentially targeted by receptor-directed ligands. Furthermore, substitutions at this conserved Gln in Gαq confer an allosteric impact different from the equivalent substitution in Gαi1, indicating distinct allosteric communication mechanisms across different Gα subtypes.

Here, we examined whether conserved residues in the Gα:Gβγ interface, in two closely related subfamilies, Gαi/o and Gαq/11, have differential effects on Gβγ coupling^12^. We chose Gαi1 and Gαq since they are representative of the two most evolutionarily related Gα subfamilies. Despite high sequence, size and structural similarity between human Gαi1 and Gαq (Figs. 1 and S1), it remains unclear whether conservation at spatially equivalent positions ensures equivalent outcomes upon mutation. The differences in their contribution to Gα:Gβγ interface coupling strength could come from spatiotemporal (spatial and time dependent) dynamic elements of the residue contacts in the interface. Therefore, we used dynamic conformational ensemble generated from all-atom Molecular Dynamics (MD) simulations of the trimeric Gαi1 and Gαq and analyzed the properties of the persistence of residue contacts, enthalpy of the interaction strength, and cooperativity using Bayesian Network Modeling (BNM), an interpretable unsupervised machine learning model. We predicted the existence of conserved residues that contribute differentially to Gαi and Gαq coupling to Gβγ and tested the predictions using bioluminescence resonance energy transfer (BRET) assays for Gβγ dissociation from the activated Gα subunit. Our results indicate that conserved positions can support subfamily-specific allosteric effects at the interface. These conserved subtype-specific hotspot residues may represent key nodes in the allosteric communication network that distinguishes Gαi1 from Gαq.

**Figure 1.**
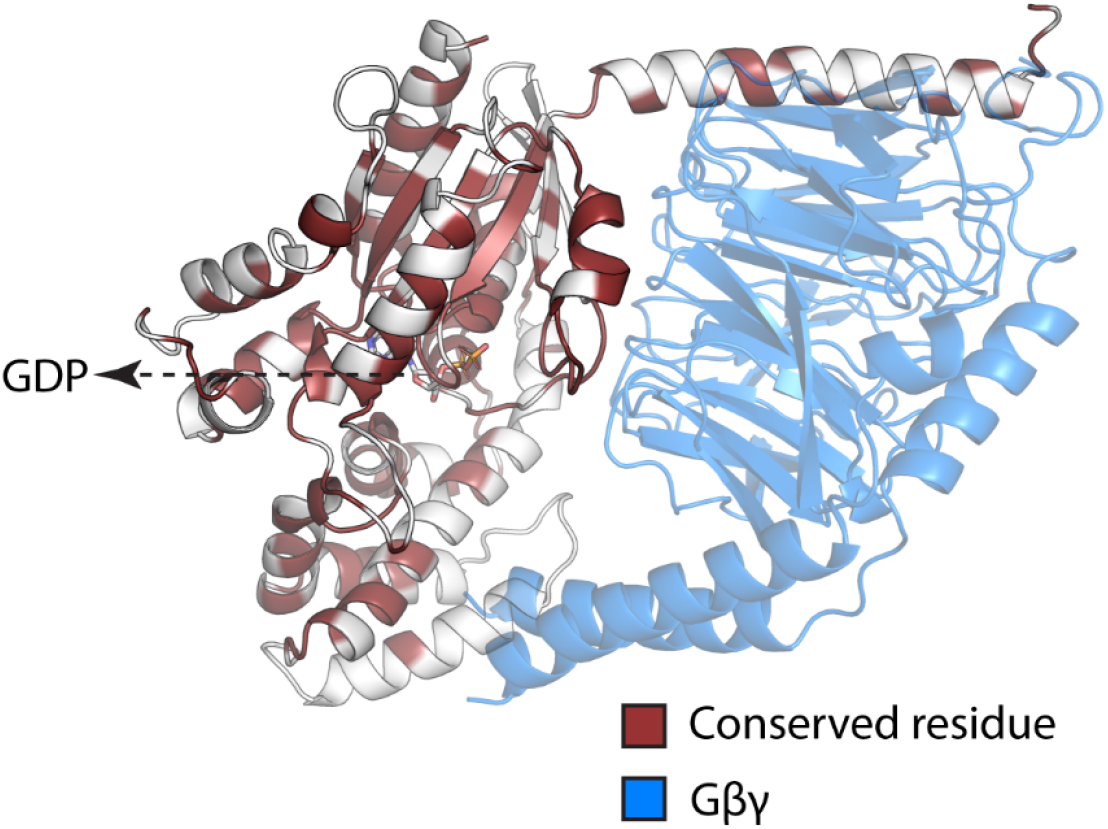
Illustration of conserved residues between Gαi1 and Gαq. GDP is shown in sticks.

## Results

### Contact frequencies and interaction energies of the interface residues show subtype-specific hotspots in the α2 helix of Gαi1 and Gαq

In this study, we investigated the role of conserved residues in the interface of Gαi1 and Gαq with Gβγ. Our goal is to identify the conserved residues in the Gα interface with Gβγ that show subtype-specific behavior. We define a hotspot as a residue that upon mutation, affects the coupling of Gα to Gβγ significantly. Furthermore, a common hotspot is defined as the residue that affects Gβγ coupling in both Gαi and Gαq subtypes equally, whereas a subtype-specific hotspot affects one Gα subtype more than the other.

We performed MD simulations on the trimeric G protein complexes starting from their respective crystal structures (Gαi1: 6CRK; Gαq: 7F6G; Fig. 2A). We performed five independent MD simulations each 1μs long, totaling 5μs for both Gαi1 and Gαq and tested each run for convergence (see Methods section). For each MD run, we calculated the contact frequencies of residue pairs in the Gα:Gβγ interface. The contact frequency is the percentage of MD snapshots that show a given residue contact in the interface. We also calculated the pair-wise interaction energies of the residue contacts in the Gα:Gβγ interface (Fig. 2A). Subsequently, we derived the per-residue contact frequencies and interaction energies for Gα residues from the above pairwise interactions (see Methods).

**Figure 2.**
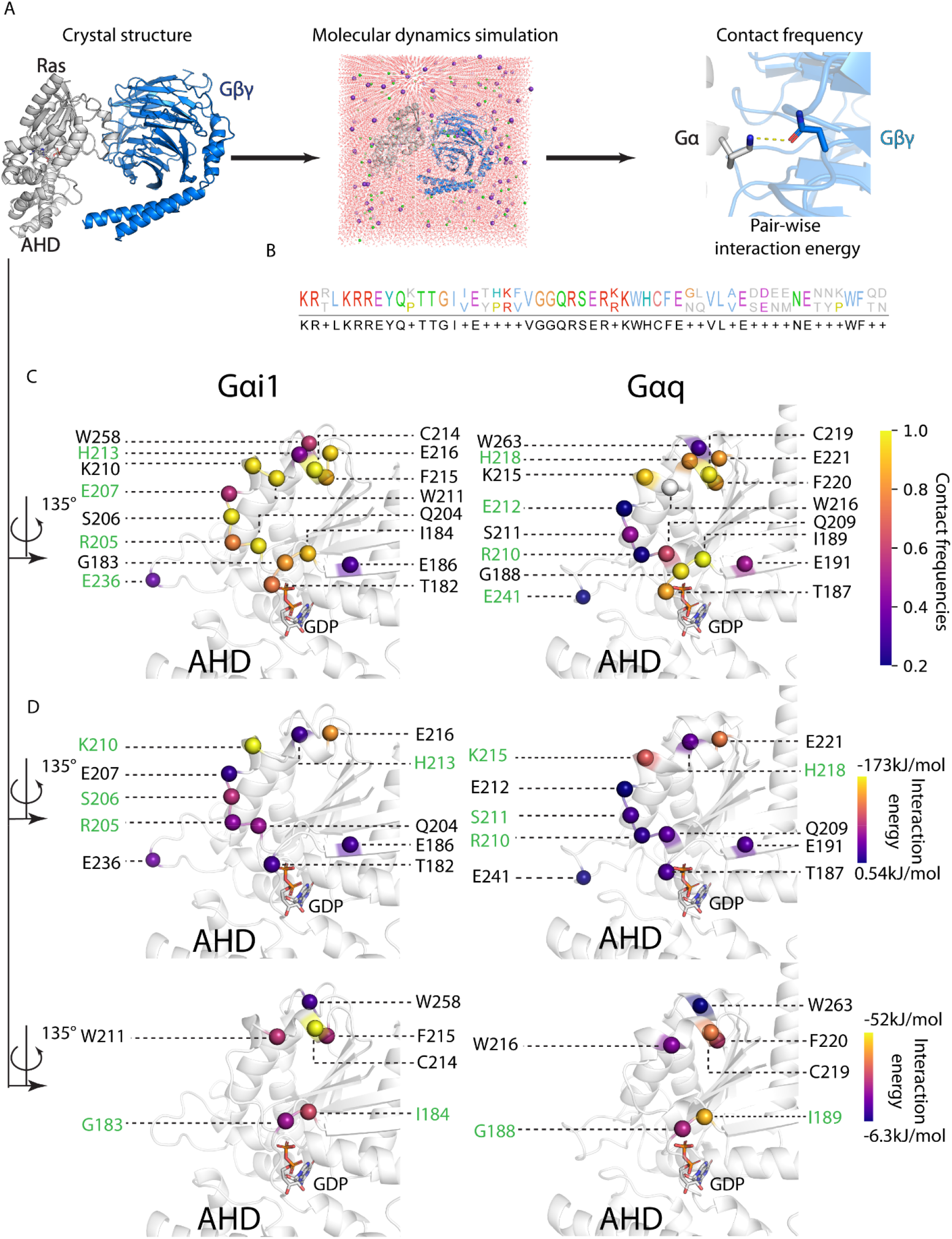
Thermodynamic properties of the interface residues in Gαi1 and Gαq with Gβγ are highly similar. **A**) Workflow showing how the MD simulations were constructed from the starting structure, and how contact frequencies and interaction energies were obtained from the trajectories. **B**) Sequence alignment of Gα interface residues. Top row is Gαi1, and bottom row is Gαq. The residues are color-coded by their physiochemical properties. Blue: hydrophobic, red: positively charged, purple: negatively charged, green: polar uncharged, yellow: cysteine, orange: glycine. **C**) Structural location of the residue contacts colored by contact frequencies of the conserved residues on the Gα subtypes. Green labels indicate residue contacts with statistical significance between the two subtypes (two-tailed Welch’s t-test, p<0.05; see Methods) **D**) Location of the residue contacts colored by their interaction energies on the interface. Top: polar residues, bottom: non-polar residues. Green labels indicate statistical significance between the two subtypes.

There are 50 amino acids in Gαi1 that contact Gβγ in the interface, and 59% of these residues in Gαi1 are strictly conserved with Gαq (Fig. 2A). There are 47 residues in Gαq that contact Gβγ in the interface, and 66% of these residues in Gαq are conserved with Gαi1 (Fig. 2A). The polar contacts are modestly enriched in Gαi1 relative to Gαq, whereas the number of non-polar residues contacting Gβγ was comparable between the subtypes.

We analyzed conserved interface residues that have contact frequencies of more than 20% (hereafter referred to as persistent contacts; Fig. S2B). Sixteen residues are conserved and made persistent contacts, out of which six are non-polar, and ten are polar. Unpaired t-test showed that 4/16 residues (25%) that show statistically significant differences in contact frequencies, are polar residues (green labeled residues in Fig. S2B). Structural mapping showed that these residues clustered predominantly within the Ras-like domain, consistent with its established importance in Gα:Gβγ interactions^1^ (Fig. 2C). Residues that show differences among subtypes are located on the α2 helix and the β4-α3 loop; these two regions are towards the periphery or outer rim of Gα:Gβγ interface. This finding is similar to our previous study showing that residues conferring Gα coupling specificity to GPCRs are also located in the outer rim of the Gα:GPCR interface^13^.

A larger set of conserved residue contacts in the Gα:Gβγ interface showed subtype-specific differences by interaction energy analysis. Because polar and nonpolar residues have distinct magnitudes of interaction energies, we analyzed them separately (Figs. 2D, S2C). 40% of the polar (Fig. S2C) and 33.3% of the non-polar interface residues (Fig. S2C) show a statistically significant difference in interaction energies. We observed two common residues (R205/R210, H213/H218) that have statistically significant differences in contact frequencies and interaction energies. Overall, measures of interaction energies revealed more differences than those of contact frequencies.

All residues that show differences in either contact frequencies or interaction energies are also highly conserved across species. There is experimental evidence that residue positions G183/G188, I184/I189, S206/S211, and E236/E241 show a significant change in coupling to Gβγ upon mutation in Gαo, a subtype closely related to Gαi1^10,14^. This data shows that contact frequency and interaction energy, when taken together, reveal subtype-specific hotspots, some of which are already reported to be mutational hotspots.

### Bayesian Network Analysis Reveals Local and Allosteric Co-dependencies in Dynamics on Gα:Gβγ interface in Gαi1 and Gαq

We have previously shown that co-dependence in the dynamic movement or correlated movement among the residues in the interface are important in protein-protein coupling and specificity ^15,16^. Residue pairs that show a high level of co-dependence in their movement with multiple residues in the interface are said to be cooperative. We next sought to identify the residue contacts in the Gα:Gβγ interface that showed cooperativity in their dynamic movement. To this end, we applied our previously developed, an interpretable and unsupervised BNM-based machine learning model, BaNDyt, to analyze MD simulation trajectories^15^. BN model analyzes the large scale data presented in the MD simulation trajectories to generate a graphical network in probabilistic space ^16^ where the nodes in the network are residues or residue contacts and the edges connecting the nodes are the probability of co-dependent motions between the nodes. Previously, we have used BN models to predict and test the amino acid residue contacts that show high cooperativity and their allosteric effect in the interface of GPCR:G protein complexes^16^. We have developed highly scalable and computationally efficient software packages for generating BN models from MD simulation trajectories: BaNDyT (specialized BN software for the MD data, freely available for download from GitHub) ^15,17–19^.

The workflow used in this study, to generate the BN models is shown in Fig. 3A (also see methods for more details). Briefly, the interaction energy of each residue in the Gα subunit within12Å of all other residues in the G protein trimer, including Gβγ residues, and nucleotide was calculated per MD frame, discretized using Max Entropy algorithm into eight bins^15.^ The resulting matrix was fed to BaNDyT software to generate the BN model. In the resulting network, the sum of all edge weights of a node (residue) is referred to as weighted degree (WD). Gα residues were ranked by their WD, and the residues in the top quartile of WD (hereafter referred to as “top residues”) were selected for further analysis. We analyzed the conserved residues in the Gα:Gβγ interface that are also in the top quartile by WD. While the number of residues in the top quartile are similar in Gαi1 and Gαq, they are not the same residues. The three topmost residues (R144, V233, and L234) are unique to Gαi1, while four other high-scoring residues (Y151, Q152, G207, and E221) are unique to Gαq (Fig. S3C). These residues are highly conserved among Gαi1 and Gαq orthologs (Table S1). Unlike subtype-specific hotspots found in contact frequencies and interaction energies, those residues have not been reported to have functional importance in Gαi1 or Gαq, except for G202/G207, of which the equivalent position was reported to be important for Gβγ subunit dissociation in Gαo ^10^.When projected to the three-dimensional structures, we observed clustering of high WD residues in the Ras-AHD interface (αD-αE loop and β4-α3 loop; Fig. 3B); these two regions separate during domain opening movements, which may confer the high co-dependencies observed.

**Figure 3.**
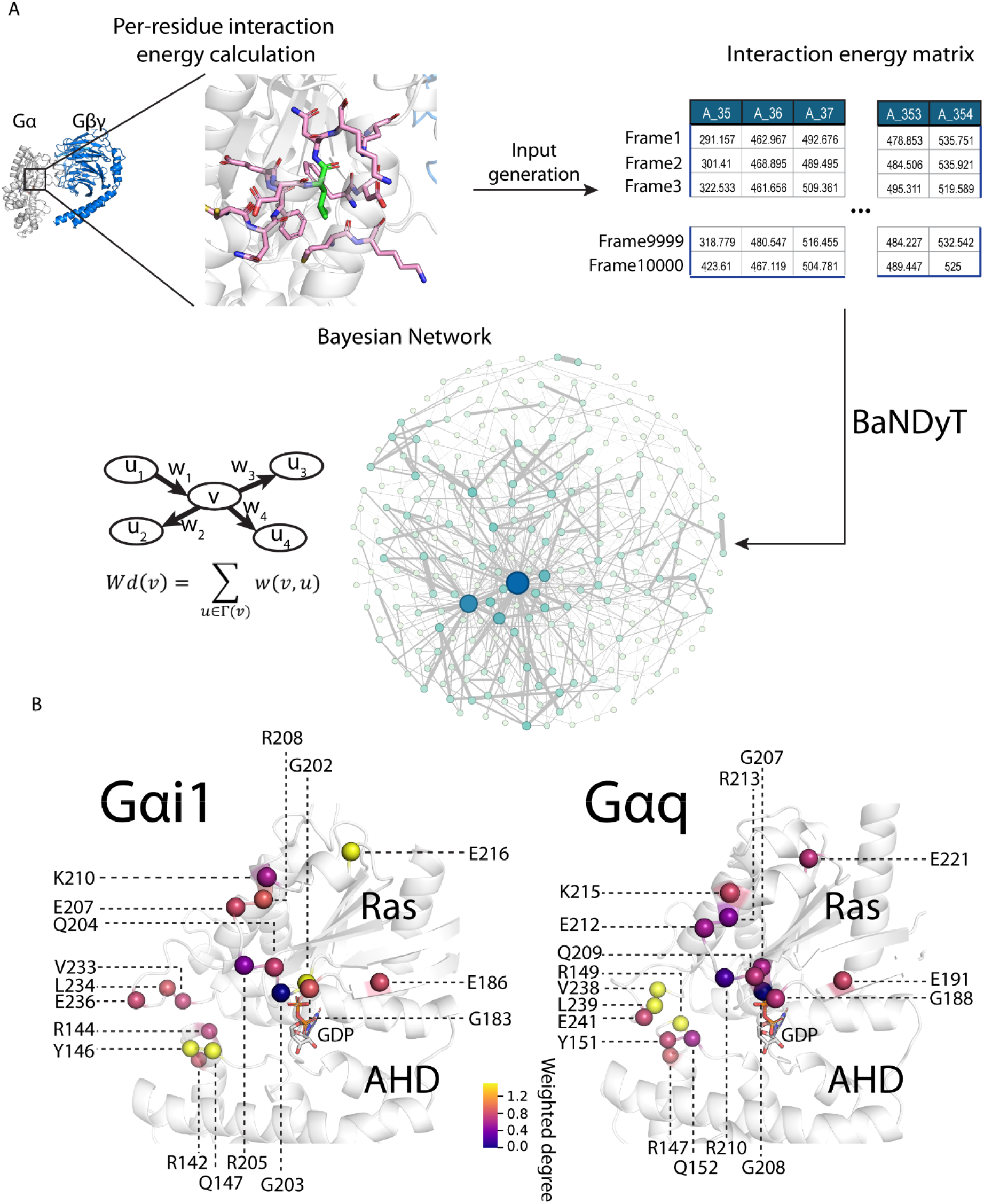
BNM shows co-dependencies in the dynamic movement among residues in the Gα:Gβγ interface and between different structural regions of Gα subunits of Gαi1 and Gαq. **A)** Schematic of the workflow used for generating the BNMs for Gαi1 and Gαq trimers. **B)** Gα-Gβγ interface residues that are also top residues projected on the three-dimensional structure. Colors of the spheres indicate the value of their weighted degrees.

Using BaNDyT, we have identified seven Gα:Gβγ interface residues that show differences in cooperativity when comparing Gαi1 and Gαq, but did not show differences in the contact frequencies and interaction energies. The lack of overlap between the subtype-specific hotspots identified using BaNDyT and those identified in residue contact frequencies and interaction energies shows that an interplay of all three factors could be important for determining the subtype specificity. Of course, other factors such as the complex cell environment are also important and not considered in this study. Here we deduced that an integration of the three features could give us a more comprehensive understanding of the intrinsic structural and dynamical features regulating the Gα:Gβγ interface.

### Distance-based prediction of residues as subtype-specific hotspots

Since no single feature was sufficient to distinguish the subtype-specific hotspot residues differentiating Gαi1 and Gαq, we inferred that subtype-specificity likely arises from a combination of the three residue-based features. Since the raw values of the three properties calculated here are of different units/dimensions we ranked the interface residues in the quantile analysis of each property to place the features on a comparable distribution. We then calculated the relative ranking of residues within each subtype (Fig. S4A). If a residue ranked the same in the two subtypes, it would have zero values for all three features and reside at the origin (Δ=0,0,0). We hypothesized that the further from the origin, the more subtype-specific a residue is in the calculated features, and the more likely that said residue will show subtype-specific effects between Gαi1 and Gαq (Fig. 4A).

**Figure 4.**
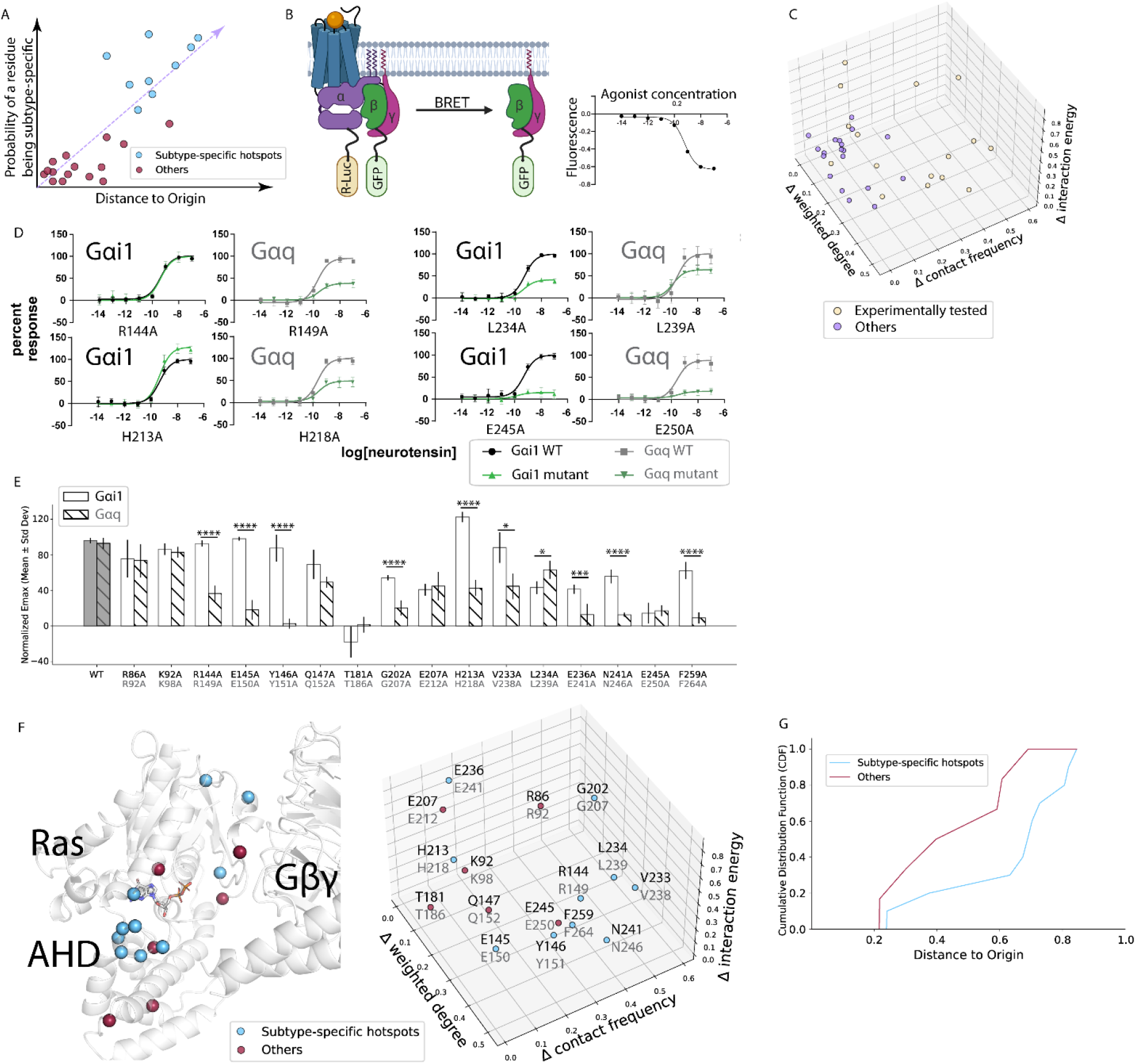
Distance-based prediction of subtype-specific hotspots and validation via TRUPATH BRET assay. **A)** Illustration of the hypothesis that the further from the origin, the more likely a residue is a subtype-specific hotspot. All points shown are conceptual and are not actual data. **B)** Schematics of the TRUPATH BRET assay. A representative conceptual readout (not actual data) is shown on the right side. **C)** Three-dimensional location of each residue with respect to the origin. **D)** TRUPATH BRET dose-response curves of representative mutants. Black/gray curve is wildtype, and green/forest curve is mutant. **E)** Normalized E_max_ values of wildtype and mutants. Stars represent statistical significance (*<0.5, ***<0.005, ****<0.001). **F)** Projection of residues on the three-dimensional protein structure (top), and annotation of tested residue in the Cartesian coordinate as seen in panel C. Cyan represents subtype-specific hotspots, Red represents other conserved residues in the interface that do not show a subtype-specific effect in the two subtypes. **G)** Cumulative distribution function (CDF) curve of residues experimentally validated as subtype-specific hotspots and those that were not (Others), as the distance to origin increases. CDFs of each group represent the accumulated proportion of values up to each distance to origin threshold, illustrating the distributional shifts between the two groups.

To validate our hypothesis, we selected 16 conserved residues along the Gα–Gβγ interface that are far from the origin, and we mutated each to alanine (Fig. 4C). BRET experiments were used to test the functional importance of these residues. Briefly, Gα and Gγ were tagged with a luciferase donor and GFP acceptor, respectively. In the inactive trimeric state, the BRET ratio is high; upon ligand-induced activation of the receptor, Gα and Gβγ dissociate and the BRET ratio decreases (Fig. 4B). When we compared the dose-response curves (Figs. 4D, S5), three distinct patterns in BRET intensity emerged: (i) subtype-specific hotspots that only affect Gαq (R144A/R149A), (ii) subtype-specific hotspots that affect both subtypes, but to a different extent (L234A/L239A, H213A/H218A), and (iii) common hotspots that affect both subtypes equally (E245A/E250A).

We observed that more mutations affected the function of Gαq than Gαi1 (Fig. 4E). Out of the sixteen residues predicted, nine were hotspots within Gαi1 (Fig. S4B), and fourteen were hotspots within Gαq (Fig. S4C). Among the sixteen tested residues, ten were subtype-specific hotspots: Gαi1/Gαq residues R144/R149, E145/E150, Y146/Y151, V233/V238 affected only Gαq, and G202/G207, H213/H218, L234/L239, E236/E241, N241/N246, F259/F264 affected both subtypes to different extent. Spatially, they are in three different structural regions of the Gα protein. G202/G207 and H213/H218 are in the interface between the Ras-like domain and Gβγ, whereas F259/F264 is located closer to Gα:GPCR interface. Most importantly, the rest of subtype-specific residues are clustered on the interface between the Ras-like domain and the AHD (Fig. 4F). This “clamp” region not only contacts the N-termini of Gβ and Gγ, but also undergoes conformational changes in the nucleotide exchange step, leading to opening of the Ras-like domain and AHD^20,21^.

To validate our hypothesis that subtype-specific hotspots are located further from the origin (Δ=0,0,0) than common hotspots, we generated a cumulative distribution function (CDF) plot (Fig. 4G). The CDF characterizes the probability that a residue assumes a value less than or equal to a specified distance from the origin. A leftward shift of the CDF curve indicates a distribution concentrated near lower distance values, reflecting a higher probability mass close to the origin. Conversely, a rightward shift signifies a distribution concentrated toward higher distance values, indicating a greater probability mass farther from the origin. As seen in Figure 4G, there is a clear separation between the two CDF curves. The CDF corresponding to subtype-specific mutants was entirely right-shifted relative to that of the common mutants, indicating consistently greater distances from the origin. This pattern suggests that subtype-specific mutations are associated with more pronounced differences in the evaluated features, while common mutations exhibit minimal variation across subtypes. The lack of overlap between the CDFs underscores a strong separation between the two distributions, therefore supporting the hypothesis. When comparing the predictive power of individual properties, we observed that contact frequencies gave the clearest separation (Fig. S4D), followed by weighted degree (Fig. S4E) and then by interaction energies (Fig. S4F). Taken together, this supported our hypothesis and demonstrated the predictive power of the three computational features.

### Allosteric communication between functionally important regions in the two Gα subtypes

The activation of the Gα subunit involves coordinated motions across structural regions that are spatially distant, and prior studies have shown the presence of allosteric communication within the Gα subunit ^1,22^. However, the mechanistic basis for allosteric communication between different structural regions of the Gα subunits, and the residues involved in such communication, are not known. Having identified conserved interface residue positions that exhibit subtype-specific impact on Gα:Gβγ dissociation, we next analyzed the mechanism of allosteric communication within the Gα protein using the BNM.

The edges in the BNM show codependency in dynamic movement between both spatially proximal residues and between allosteric or spatially distant residues. We calculated allosteric co-dependencies (edge strength) from BNM to identify distant residue pairs that show long range correlated movement. We summed up the edge strengths of all edges that connect residues from two structural regions to uncover how different structural regions show correlated movements. This would lead to an understanding of the allosteric communication mechanism within the Gαi1 and Gαq subtypes. We constructed radial graphs (also known as circos maps) shown in Supplementary Figure S6 that show the total edge strengths between structural regions connecting local and distant residues in Gαi1 and Gαq. For both Gα subtypes studied here, we observed strong correlated motion or co-dependencies in the BNM between residues in the structurally neighboring regions, indicating impact of movement of neighboring residues in a local environment. Structural regions containing Gα:Gβγ interface residues (αD-αE loop, α2 helix, β4-α3 loop) showed correlated movement with distant (allosteric) residues within the interface. Furthermore, they also showed correlated movement to structural regions beyond the Gα:Gβγ interface, namely α1-β1 loop, α1-αA loop, αA helix, and β5-αG loop, all of which are distant from the Gα:Gβγ interface. Interestingly, while α5 of Gαi1 showed allosteric correlated movement only with other Ras-like domain regions (α2, α3, and α4-β6 loop), α5 of Gαq also had allosteric correlated movement with AHD regions (α2, αC, αF). This is potentially significant given that the α5 helix extends into the receptor cavity and represents a major determinant of Gα coupling specificity. We also observed correlated movement between nucleotide binding site regions and the Gα-Gβγ interface (i.e. between β1-α1 loop and β4-α3 loop), as affirmed by our previous study^1^.

Having considered allosteric communication with receptors and nucleotides, we examined the existence of allosteric communication or correlated movement in regions where the Gα subunits bind to effectors and regulators. We constructed a multi-scale meta-network representation of BNM in Gα. We first delineated the Gα residues located in the interaction interfaces of Gα with its well characterized binding partner proteins such as: the GPCRs, Gβγ, Regulator of G-protein Signaling (RGS), GDP-binding pocket, and Ric8 guanine-nucleotide exchange factor (see Methods, Fig. 5A). The residues of Gα involved in these interfaces were calculated using GetContacts from the respective crystal structures (See Methods, Tables S2, S3). Each “meta-node” corresponds to one of these interfaces (Fig. 5B), and each edge in the meta-network encodes the correlated motions calculated between pairs of interface residues (sum of all edge strengths). The edge weights in the meta-network were computed as the sum of all edge weights connecting residues belonging to the corresponding interface residue pairs. Weighted degrees of meta-nodes were obtained by summing the edge strengths of all meta-edges incident on that node. This abstraction enabled a compact yet informative summary of dependencies that link functionally active regions within each Gα subunit.

**Figure 5.**
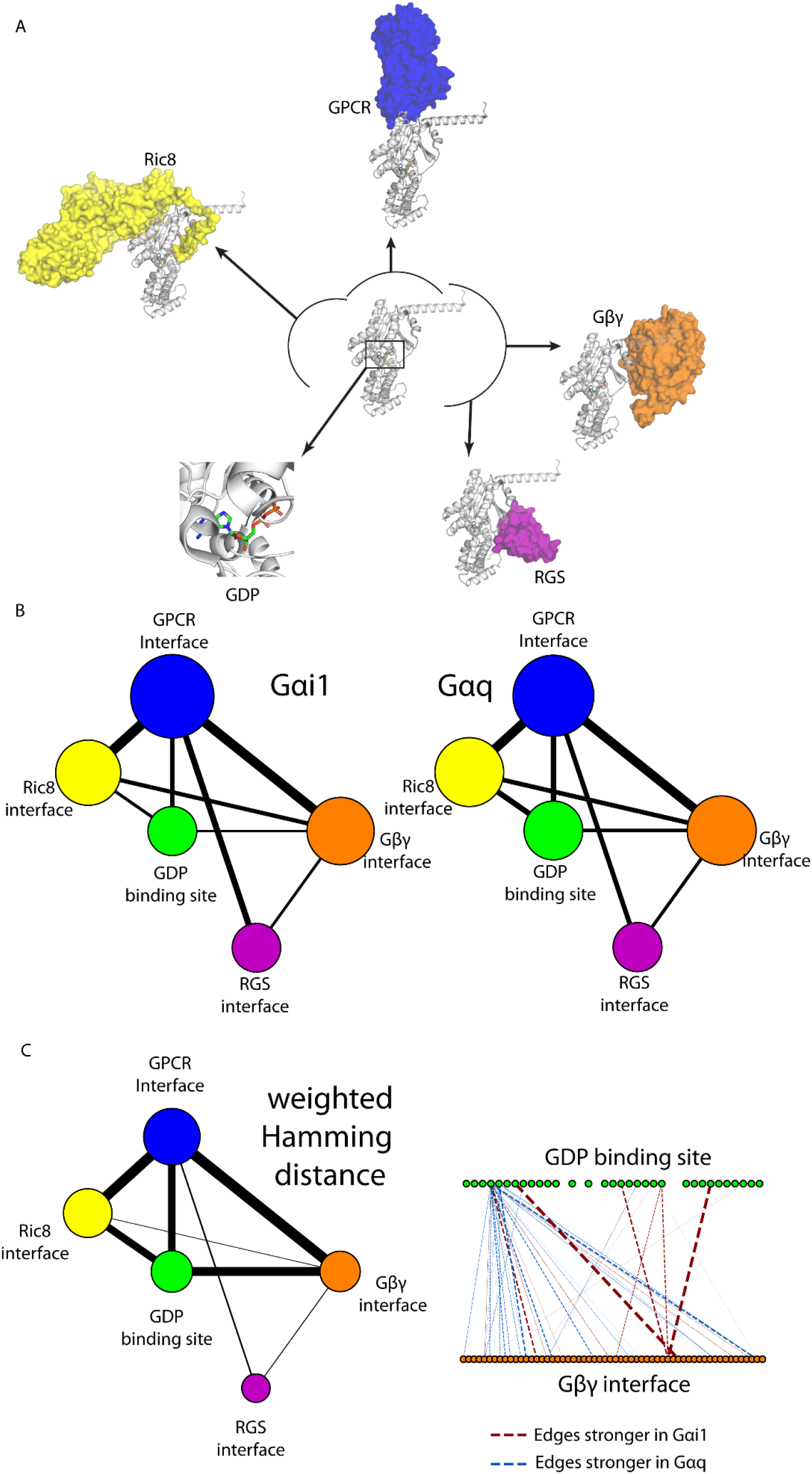
BNMs show the Allosteric Communication between different protein interaction interface sites in Gα subunits. **A)** Interface locations of proteins that are known to interact with the Gα subunit. In a clockwise direction from top: G protein-coupled receptor (GPCR), Adenyl Cyclase, Gβγ dimer, Regulator of G protein Signaling (RGS), GDP, and guanine nucleotide exchange factor (Ric8). **B)** Meta network illustrating the correlated movements between the interfaces to the major interacting partners. Blue: GPCR interface, Orange: Gβγ interface, Purple: RGS interface, White: GDP binding site, Yellow: Ric8 interface. The sizes of the meta nodes represent their weighted degrees, while the width of the meta edges represent their weights. C) Weighted Hamming distance network (left) and edge distance break down between GDP binding site and Gβγ interface (right). Maroon: edges that are stronger in Gαi1; marine: edges that are stronger in Gαq.

The resulting meta-network, or higher order structural level network, models for Gαi1 and Gαq shown in Figure 5B displayed strikingly similar global topologies. In both subtypes, we observed strong correlated motions between the residues in the GPCR:Gα interface and those in Gα:Gβγ interface, followed by GPCR:Gα interface to the Ric8 interface and to the RGS interface. GPCRs and Ric8 function as guanine nucleotide exchange factors (GEFs) for Gα subunits ^23^, while RGS proteins act as GTPase-activating proteins (GAPs) ^24^, collectively regulating the GTPase on-off cycle. The observed correlated movements among the GPCR, Ric8, and RGS interfaces suggest that these regulatory sites are intrinsically coupled within the conformational landscape of the Gα subunit. Such coordinated dynamics likely reflect allosteric networks that facilitate transitions through nucleotide exchange and hydrolysis, even in the absence of bound effectors.

Although the overall meta-network between the different structural regions look similar in Gαq and Gαi (Fig. 5B) we observed changes in the edge strengths between the residues in the different structural regions. To identify these changes, we calculated the weighted Hamming distance that quantifies differences between corresponding edge strengths in Gαi1 and Gαo (Fig. 5C, left). Indeed, while the weighted degree difference of Gβγ interface residues is very low (0.4 bit between Gαi1 and Gαq networks), the weighted Hamming distance is much higher (24.0 bits between Gαi1 and Gαq networks). The discrepancy between the values suggests that while the importance of Gβγ interface residues is comparable in Gαi1 and Gαq, they exert their influences through distinct routes. To illustrate the idea, we compared edge weights connecting GDP binding site and Gβγ interface residues (Fig. 5C, right). We observed that while most of the edges is stronger in Gαq, a few edges are much stronger in Gαi1. The more dispersed connectivity found in Gαq could explain the general sensitivity of Gαq to mutations: out of the 16 proposed mutations, 14 affected the dissociation of Gαq.

## Discussion

Although conserved amino acids at the interface of proteins are assumed to have similar effect, our study revealed that certain conserved amino acids on Gαq and Gαi1 subtypes show differential effects on coupling to the Gβγ subunits. These differences primarily stem from the dynamics of the Gα subunit. By integrating three complementary features derived from MD simulations, namely: (i) residue contact frequencies, (ii) residue interaction energies, and (iii) co-dependencies among the residue pairs derived from Bayesian Network Analysis, we showed that the subtype selective coupling of Gαi1 and Gαq comes from a complex confluence of temporal persistence of residue contacts, as well as from enthalpic and cooperative effects. This triangulation revealed subtype-specific interface characteristics that become apparent only from the dynamic ensemble analysis. Through an iteration of prediction and experimental testing we have identified conserved amino acid residues in Gαq and Gαi1 that have subtype-specific effects on coupling to Gβγ subunits. This supports the hypothesis that conserved residue positions can elicit distinct functional outcomes in paralogous proteins.

Switch II (α2 helix) and switch III (α3 helix) make direct contacts with Gβγ and undergo substantial conformational changes upon Gα activation. Therefore, residues within these two regions have been the central focus of prior studies investigating determinants of Gβγ release. Consistent with previous studies,^1,10^ mutations at Switch II and III were disruptive in both Gα subtypes in our study. Beyond the two canonical regions, our approach also highlighted previously understudied structural regions of the AHD and identified hotspot residues that affect Gβγ coupling upon mutation in Gαq but not in Gαi1. These observations suggest that the AHD, a region primarily studied for its interaction with RGS and its function in facilitating nucleotide exchange and hydrolysis ^25–27^, may play a pivotal role in Gβγ dissociation. Those hotspots in the AHD along with a few others in the Ras-like domain are clustered around the Ras-AHD clamp region. This clamp region mediates the opening and closing motion of the Ras domain and AHD that is necessary for nucleotide exchange during Gα activation. We speculate that these differences in the clamp regions can lead to differences in nucleotide exchange rates ^20,28,29^. The subtype dependent differences we discovered in this region are consistent with previously reported differences in Gβγ release kinetics^30^. However, it should be noted that we did not calculate any kinetic properties of Gβγ dissociation from our MD simulations. The recovery of previously reported hotspots, together with the discovery of new subtype-specific hotspots, provides internal support for our computational pipeline, which combines classical physical interaction metrics with a probabilistic machine learning network model to estimate cooperativity in coupling^1,10^.

Our computational features capture both the effect of local environments and allosteric communication of the interface residues. A previous study by Hewitt-Valentin et al has shown that the conserved “catalytic Gln” residue that is outside the Gα:Gβγ interface showed subtype-specific effects on Gαq and Gαi1, and these effects were the result of long-range allostery. The present study focuses on conserved residues within the interface because these positions provide the most challenging readouts of whether spatially equivalent conserved residues produce equivalent perturbation outcomes across paralogs. Building on our new and previous work^31^, future studies will examine the role of conserved residues outside the Gα:Gβγ interface and their allosteric contributions to differential Gα:Gβγ coupling, thereby extending the scope beyond the central questions addressed here.

More broadly, these findings indicate that paralogous signaling proteins may harbor subtype-specific mutational sensitivities even when their sequences and global structures are largely conserved. Our strategy of integrating interface energetics, cooperative coupling, and mutational assessment is broadly applicable to other paralogous proteins. As efforts toward precision pharmacology increasingly consider G-protein subtype selectivity, analytical approaches capable of identifying non-obvious determinants of specificity may prove valuable for future therapeutic design.

## Methods

### Multiple sequence alignment

The sequences of human Gαi1 and Gαq were retrieved from UniProt (https://www.uniprot.org/) with accession code P63096 and P50148, respectively. The alignment was performed using ClustalW web service (https://www.genome.jp/tools-bin/clustalw) and visualized using Jalview^32^. For the alignment of the interface residues, the interface residues were extracted from the contact results and aligned using the same procedure as the full sequence alignment.

### Details of Molecular Dynamics (MD) simulations

The structure of GDP-bound Gαi1-Gβγ complex was downloaded from RCSB (PDB ID: 6CRK). To generate a structural model for the GDP bound inactive state of Gαq:Gβγ complex, we downloaded the structure of apo Gαq-Gβγ complex from RCSB (PDB ID: 7F6G), in which the AHD of Gαq was focus resolved. GDP was transferred to this Gαq structure from 6CRK, by aligning Gαi1 and Gαq by their Ras-like domain. The αN helix was removed from both structures; we performed the simulations in soluble state, and without membrane anchoring the αN helix would become unstable. We optimized the side chain rotamer conformations of the residues within 7Å of GDP, followed by energy minimization using conjugate gradient method with a convergence cutoff of 0.1kcal/mol Å^2^. Both operations were performed using the Prime module of Maestro software from Schrodinger (https://www.schrodinger.com/products/maestro). Input structure files for MD simulations were generated using CHARMM-GUI^33^. Each complex was solvated in explicit TIP3P water molecules in a cubic box (11.5nm x 11.5nm x 11.5nm) with 0.15M of potassium chloride for maintaining the physiological condition. We used software GROMACS^34^ (Version 2022.3) with all-atom CHARMM36m force field^35^ to perform MD simulations at 310°K coupled to a temperature bath with a relaxation time of 0.1ps^36^. Desired pressure for all systems were achieved by using Parrinello-Rahman barostat with a pressure relaxation time of 2ps^37^ Equilibrium bond length and geometry of water molecules were constrained using the SHAKE algorithm^38^. The short-range electrostatic and van der Waals interactions were estimated every 2fs timestep using a charged group pair list with cutoff of 8Å between centers of geometry of charged groups. Long-range van der Waals interactions were calculated using a cutoff of 14Å and long-range electrostatic interactions were treated with the particle mesh Ewald method^39^. Temperature was kept constant at 310°K by applying the Nose-Hoover thermostat^40^. Before production runs, we equilibrated the systems using the following procedure: All systems were subjected to a 5000-step steepest descent energy minimization to remove clashes^41^. After minimization, the systems were heated up to 310°K under constant temperature-volume ensemble (NVT). The simulations were saved every 200ps for analysis. The protein and nucleotide were subjected to positional constraints under a harmonic force constant of 1000 kJ/(mol*nm^2^) during the NVT step while solvent molecules were free to move. The systems then were further equilibrated using constant pressure ensemble (NPT), in which the force constant applied to the protein, Mg ion, and nucleotide were gradually reduced from 5kJ/(mol*nm^2^) to zero in six steps of 5ns each. An additional 50ns of unconstrained simulation was performed, making it a total of 80ns NPT equilibration prior to production runs. We performed five production runs of 1000ns each using five different initial velocities for every system. We collected the five trajectories to obtain a 5μs MD trajectory for both Gαi1-Gβγ complex and Gαq-Gβγ complex. The convergence of each MD trajectory was validated by plotting the time series of the RMSD in the coordinates of the backbone atoms of the G protein trimer with respect to the starting structure of production run (Fig. S7). All the runs show a RMSD converging at around 7 Å and most of these RMSD come from the relative movement of the Ras-like domain and AHD and movements of Gβγ N termini.

### Per-residue contact frequency of Gα residues

For each system, the last 600ns from five independent MD simulation runs were merged into one concatenated trajectory. The concatenated trajectory was sampled every 2ns. The sampled trajectory was fed to GetContacts script available on GitHub (https://github.com/getcontacts/getcontacts). For specific parameters, the interaction type flag “itype” was set to “all”, and the two atom group flags for contacts were “chain A” and “chain B or chain G”. Contact results were further processed in Python environment: briefly, the contact frequency of a Gα residue is calculated as the percentage of MD frames in which said residue makes any contact with any residue on Gβ protein or Gγ protein. The contact frequencies were used to generate Fig. 2C. Residues with contact frequencies greater than 20% were used for plotting Fig. S2B.

### Per-residue interaction energy of Gα residues

For each system, we aggregated the last 600ns of each trajectory run to calculate the pair-wise interaction energies of contact pairs between Gα and Gβγ, sampled every 2ns. For analyzing the interface residues, each residue of a contact pair was grouped into an energy group in GROMACS, and total non-bond interaction energy (van der Waals + Columbic) was calculated using gmx energy module. The per-residue interaction energy of a residue was calculated by summing the interaction energies of all contact pairs said residue was involved in.

### Bayesian Network Model Generation

The BNM was constructed using BaNDyT ^15^ software developed by us, using the workflow shown in Fig. 3A. The input feature to generate the BNM was interaction energy of each residue with all the residues within 12Å in each MD snapshot. For each G protein system, we aggregated the last 600ns of each of the five trajectories to calculate the pair-wise interaction energies of contact pairs between Gα and Gβγ, sampled every 2ns and calculated the per residue interaction energy for each MD snapshot. The interaction energies were assembled in a matrix with each row representing a frame from the trajectory, and each column representing a residue (Fig. 3A). The two matrices acquired from MD trajectories of Gαi1 and Gαq were stacked vertically, keeping only the shared columns. The distribution of the interaction energies for each residue across the MD simulation trajectories given in the concatenated matrix was discretized using MaxEnt into eight bins. This was fed to the BANDyT workflow to generate a Bayesian network universal graph^42^. Once the universal graph was recovered, the individual networks were obtained by rescoring against stratified data, using mutual information between the nodes. The output was further processed in Python environment.

The network was visualized using Gephi (https://github.com/gephi). The network was saved in graphml format and fed to Gephi. The weighted degree for each node (residue in Gα subunit) was calculated as the sum of all the edge strengths connected to the given node (see Fig. 3A). The color and the size of the nodes are proportional to the weighted degree; the width and color of the edges were proportional to the edge weight.

For meta-network generation, the residue-level network was coarse-grained by summing residues into groups (Gα–Gβγ interface, GDP-binding site, GPCR interface, and Ric8 interface). The interfaces were defined as follows: Gα–Gβγ interface was defined by GetContacts, described above. GDP-binding site was defined by GetContacts; specifically, the topology and trajectory used were the same as described in section “*Per-residue contact frequency of G*α *residues*”, the group selection used were “resname GDP” and “chain A”, with “itypes” flag set to “all”. The definition of GPCR interface, adenyl cyclase interface, RGS interface, and Ric8 interface was taken from a previous publication^43^. Briefly, the protein entity information was retrieved from RCSB (www.rcsb.org), and GetContacts were used to calculate the interacting residues between Gα subunit and its partner proteins. The interface residues can be found in Table S2. The PDB structures used can be found in Table S3. Each structural region forms a meta-node, and the meta-edges represent the summed pairwise edge weights between residues in the corresponding regions. Meta-node size is proportional to residue count, and meta-edge width corresponds to the aggregated inter-region connectivity. Ras-other and AHD-other encompass residues that are in Ras and AHD, respectively, but are not part of the functionally defining regions.

### Robustness test for Bayesian Network Model

The concatenated trajectory comprised 10,000 frames organized into five independent segments of 2,000 frames, each corresponding to an independently initialized velocity. Bootstrap resampling was performed with replacement within each segment, preserving the original segmentation. One thousand bootstrap ensembles were generated. For each ensemble, network edges were re-estimated and scored against the universal graph using mutual information between node pairs. The confidence interval was calculated as an evaluation. The relevant data can be found in the supplementary information matrix 1 (weighted degree edge confidence interval) and matrix 2 (weighted hamming distance edge confidence interval).

### Bioluminescence resonance energy transfer (TRUPATH BRET^44^)

HEK293T/17 cells (ATCC; CRL-11268) were maintained at 37°C, 5% CO_2_ in a HERAcell VIOS 160i CO_2_ incubator (Thermo Fisher Scientific; 51033547) in Dulbecco’s Modified Eagle Medium (DMEM)(Corning; 10013CV) with 10% fetal bovine serum (Sigma-Aldrich; F2442), 100 U/mL penicillin, and 100 μg/mL streptomycin (Gibco; 15140122). 5-8 hours after seeding (1,000,000 cells per 3 mL), the cells were transfected with 300 ng human neurotensin receptor 1 (NTS1R), Renilla luciferase 8 (Rluc8) tagged human G i1 or G q, human Gβ3, and green fluorescent protein 2 (GFP2) tagged human Gγ9 in a 1:1:1:1 ratio using TransIT®-2020 transfection reagent (MirusBio; MIR 5404).1 14-18h post transfection, the cells were seeded on a 96-well plate (Corning; ref 3903) coated with poly-D-lysine (Gibco; A38904-01) at a density of 80,000 cells in 150 µL 1% FBS and 1% PEN/STREP DMEM per well. The next day, the plates were washed with 60 μL per well of drug buffer (20 mM 4-(2-hydroxyethyl)-1-piperazineethanesulfonic acid (HEPES) (Thermo Fisher Scientific; J16924.K2), 0.3% bovine serum albumin (Sigma-Aldrich; A6003), 0.03% L-ascorbic acid (Sigma-Aldrich; A92902), in Hanks’ Balanced Salt Solution (Gibco; 14025092), pH 7.4) for 10 min at 37°C before the addition of Rluc8 substrate, coelentrazine 400a (NanoLight; 340), at 7.5 µM (1.5x) in 60 µL drug buffer. After a 10 min incubation in the dark at room temperature, 30 µL (3x) of neurotensin (Tocris; 1909) in drug buffer was added to the plate for final well concentrations ranging from 100 µM to 0 µM. Following a 10 min dark incubation, the plate luminescence was read by a CLARIOstar Plus (BMG LABTECH) in a 4 mm spiral well scan at 410 nm (Rluc8 bioluminescence) and 530 nm (GFP2 fluorescence). The BRET (GFP2 emission/ Rluc8 emission) ratio was baseline corrected in GraphPad Prism 10 (GraphPad Software Inc., San Diego, CA) by subtracting the respective zero agonist BRET ratio from each sample to yield net BRET. Net BRET values were then normalized to the response of their respective wildtype (G i1 or G q) to provide percent response of each mutant.

The basal BRET values of mutants were normalized to their respective wild-type, and plotted in Fig. S8.

The dose-response curves were plotted in GraphPad Prism v10.6.1. The curves were generated by non-linear fit using log(agonist) vs. response mode. Coloring and labelling of the curves were done in Adobe Illustrator.

### Figure generation of computational data

Heatmap was generated using Seaborn package in Python. We used the color map “plasma”, with the lower bound set to 20% for contact, and the maximum value for interaction energy and the upper bound set to 100% for contact, and the minimum value for interaction energy. The labels of heatmaps were generated in Adobe Illustrator.

PYMOL figures were generated using the plotting array of the heatmap. The Cα atoms of the residues were colored according to their values. We adopted the same color map as the heatmap. The nucleotide was colored per atom type, with carbon set to white (pymol.cmd.util.cbaw).

The circos maps of the BNM were generated using Python Holoview package. The nodes were ordered according to the sequence and labeled using their region names. The labels were further processed in Adobe Illustrator.

### Statistical analysis

For contact frequency and interaction energy, the values from each frame was compiled as an array, and two-tailed Welch’s t-test was applied to calculate the p-value. P-values less than 0.05 was considered statistically significant.

## Supporting information

Supplementary information matrix 1

Supplementary information matrix 2

## Data Availability

The MD trajectories are available on GPCRmd.org. They will be made available upon acceptance.

## Supporting Information

This article contains supporting information.

## Funding and additional information

Funding for this work was supported by NIH grant R35 GM156498 to N.V., 5R35 GM118105 to H.G.D, R01-LM013876 to N.V., A.S.R and S.B, and R01-LM013138 to A.S.R. Additional support is acknowledged by Dr. Susumu Ohno Endowed Chair in Theoretical Biology (held by A.S.R.)

