## Supplementary information for "Conserved Residues in the Gα interface show subtype specificity in Gβγ coupling"

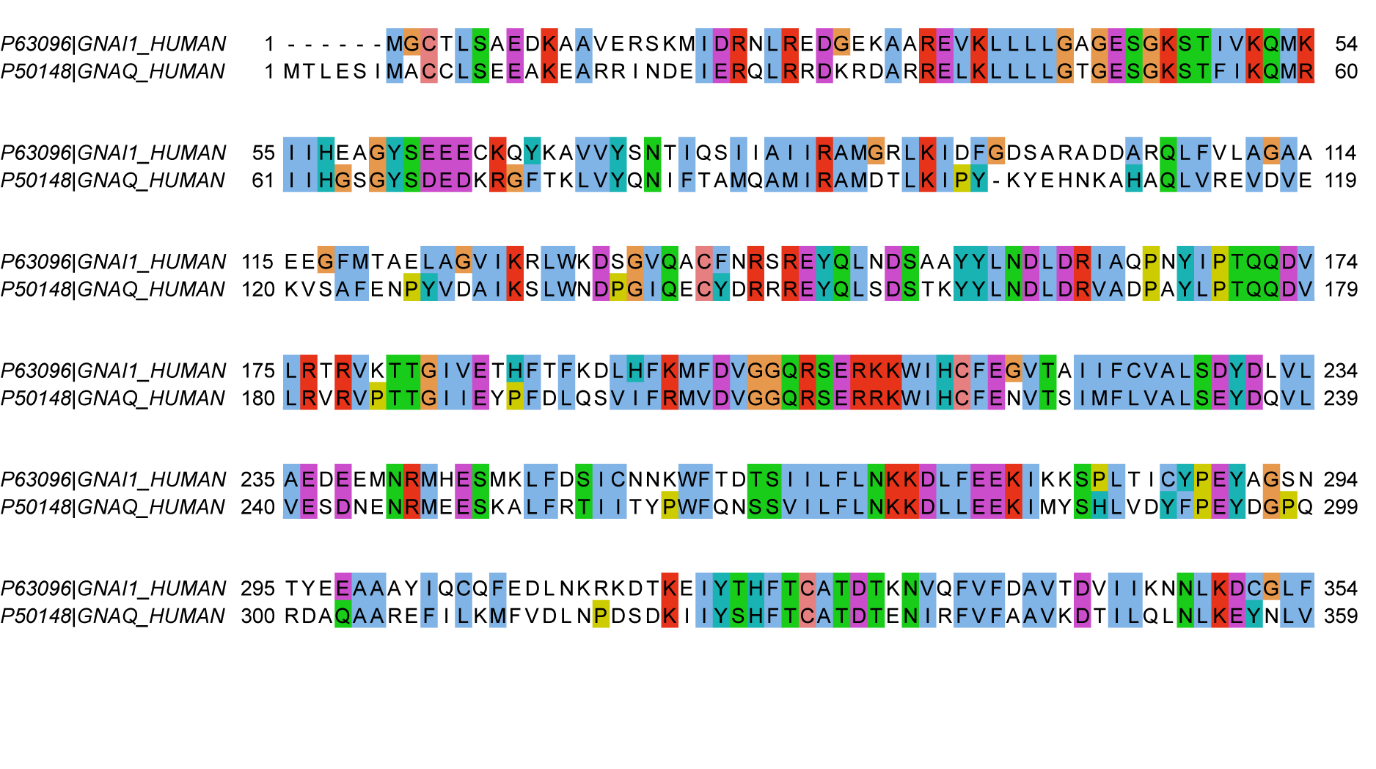
**Figure S1**. Sequence alignment of human Gαi1 and human Gαq. The residues are color-coded by their physiochemical properties. Blue: hydrophobic, red: positively charged, purple: negatively charged, green: polar uncharged, yellow: cysteine, orange: glycine.

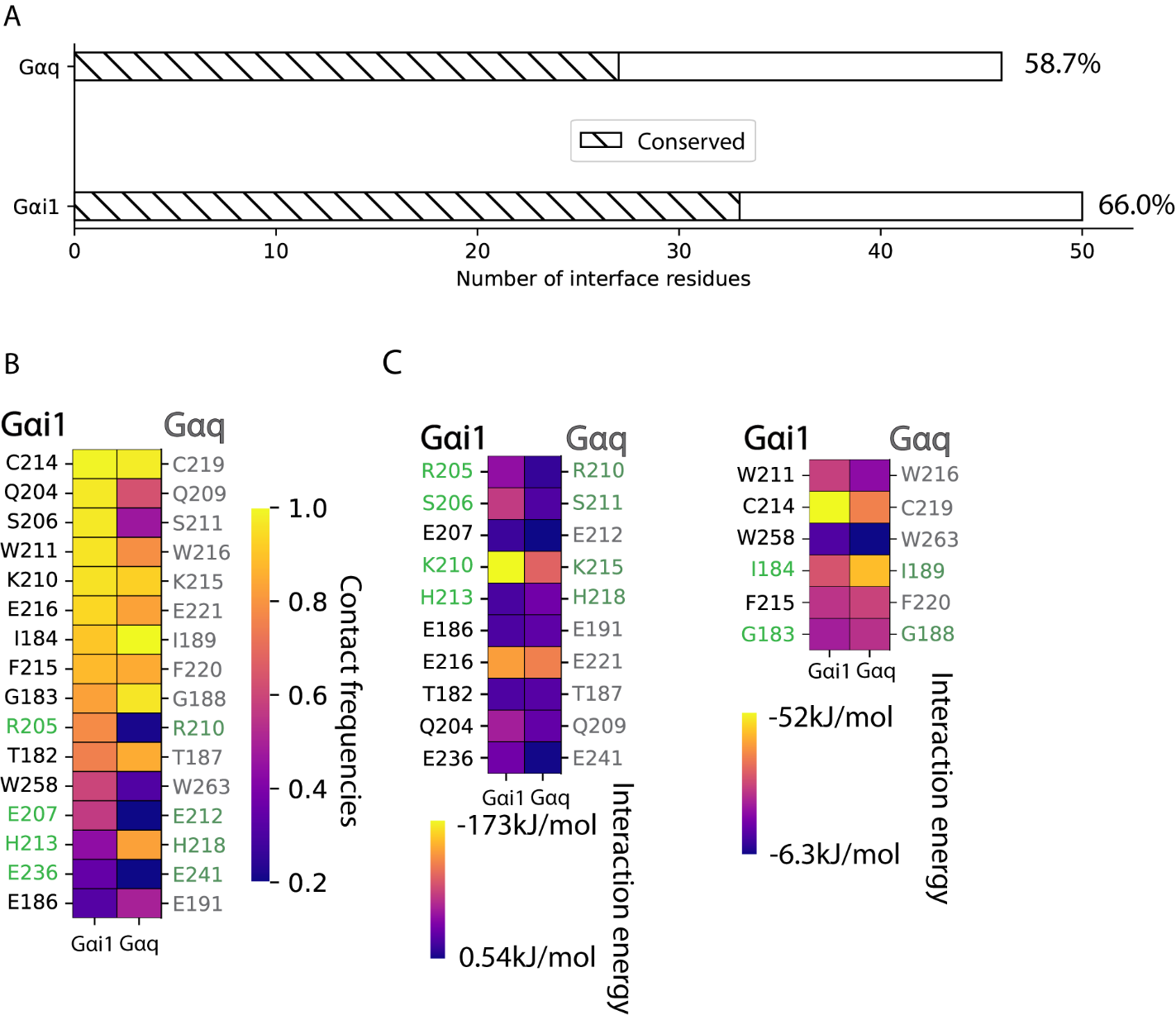

**Figure S2**. Percentage sequence conservation and thermodynamic properties of Gα-Gβγ interface. A) The number of conserved residues among all the interface residues. Percentages of conserved residues are annotated in values next to the bar. B) Heatmap of contact frequencies of interface residues. Green labels indicate residues that show statistical significance in the respective properties. C) Heatmaps of interaction energies of interface residues. Left: polar residues, right: non-polar residues. Green labels indicate statistical significance.

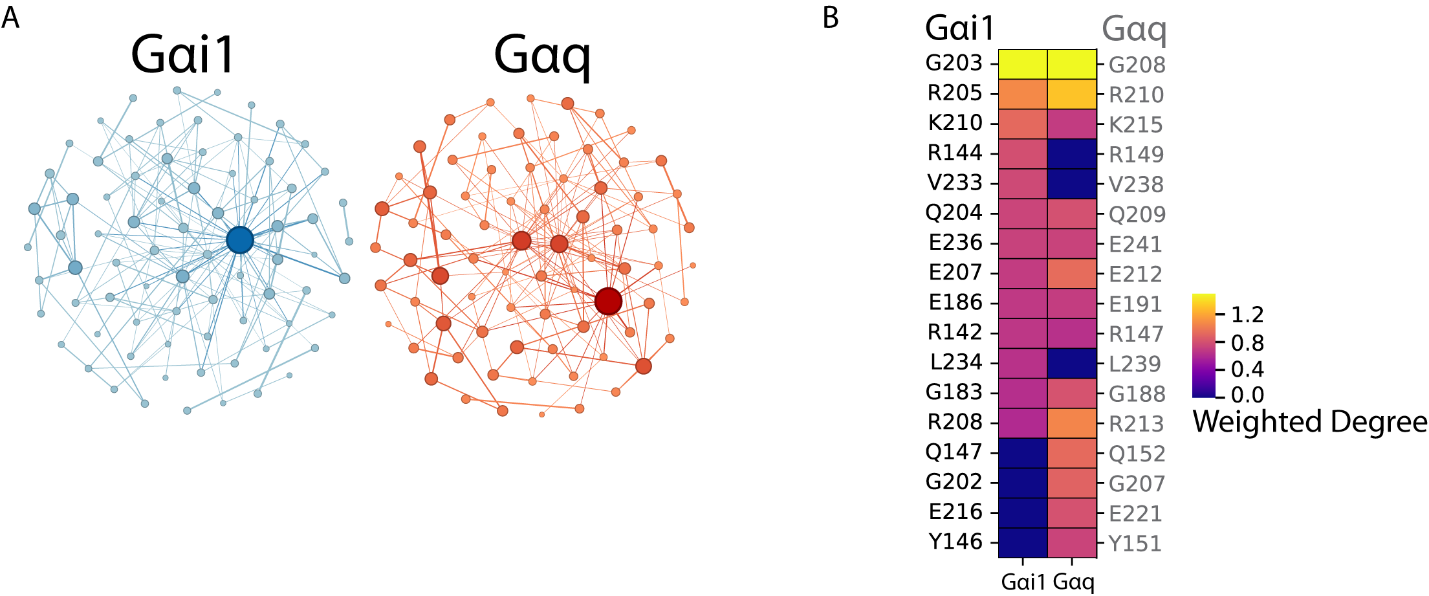

**Figure S3**. Co-dependency matrix visualization and breakdown. A) Top scoring residues networks of Gαi1 and Gαq. B) Heatmap of weighted degrees of Gα:Gβγ interface residues.

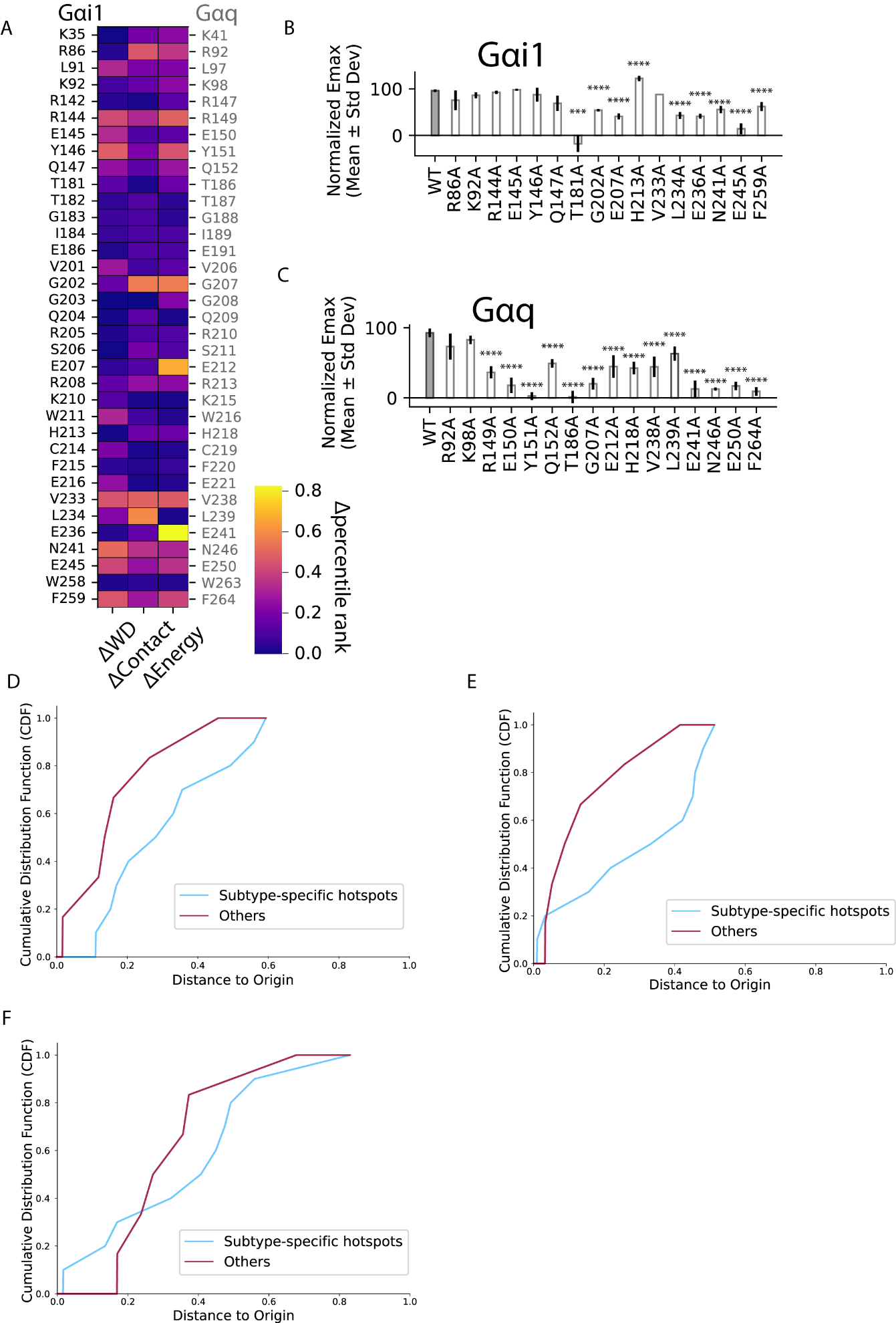

**Figure S4**. Raw data of distance calculation, structural location of tested residues, and raw BRET data. **A)** Compiled data for distance calculation. The values represented were percentile rankings among all interface residues. **B & C**) Comparison of BRET signals of mutants to their respective wildtypes. **D, E, F**) Cumulative distribution function (CDF) plot of contact frequencies, weighted degree, and interaction energies respectively. The CDF curves show the proportion of residues with values less than or equal to a given distance. It helps visualize how the properties are distributed across the two systems.

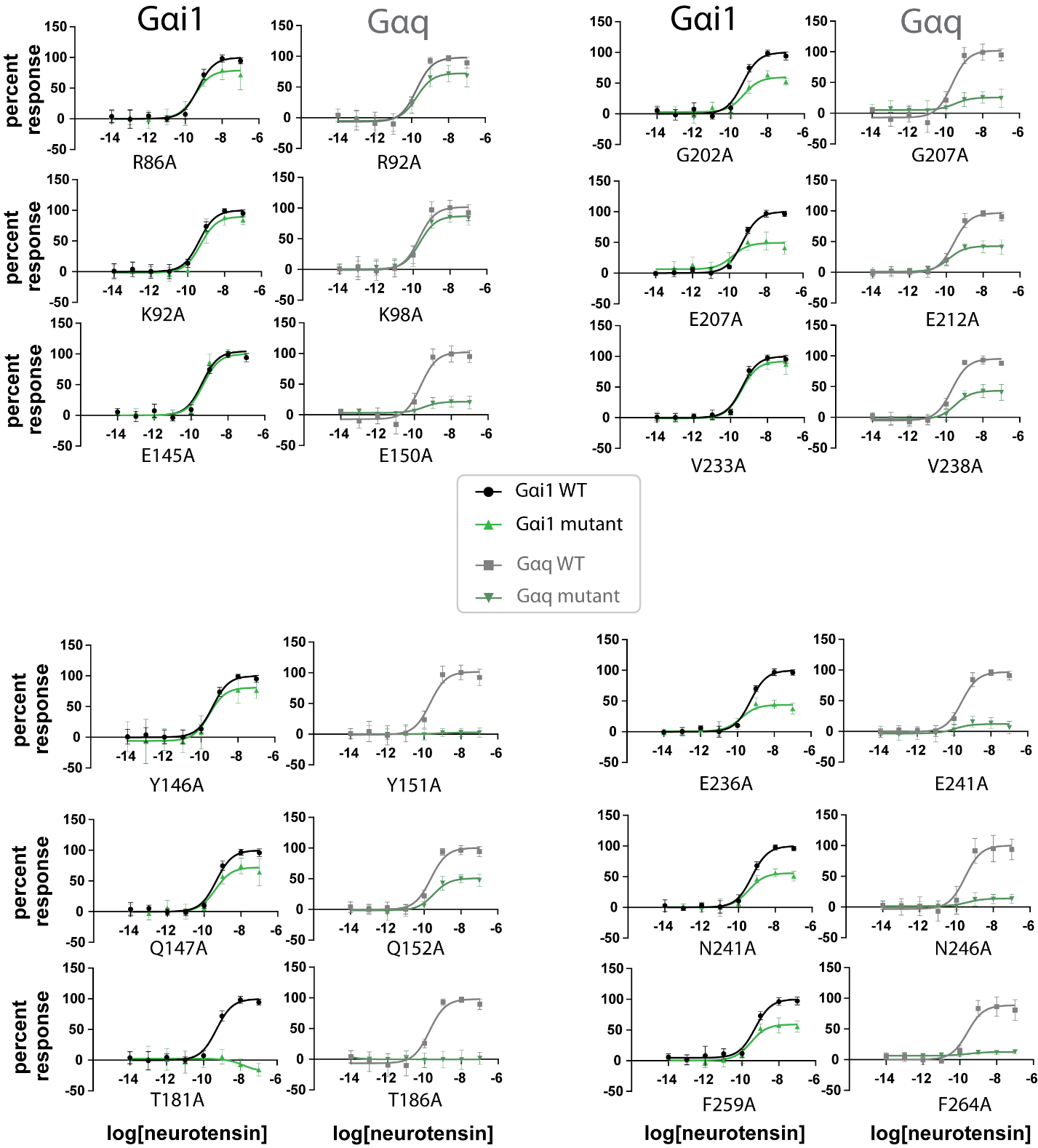

**Figure S5**. Dose-response curves of BRET signaling in response to increasing concentrations of the GPCR agonist neurotensin. All conditions were done in triplicates, and the dots showed the average. Y-axis is the percentage of BRET signaling loss in comparison to the average values of wild-type, representing the percentage of dissociated Gβγ upon GPCR-mediated Gα activation. The X-axis is the log-normalized concentration of neurotensin. Black: Gαi1 wild-type, gray: Gαi1 mutants, green: Gαq wild-type, forest: Gαq mutants.

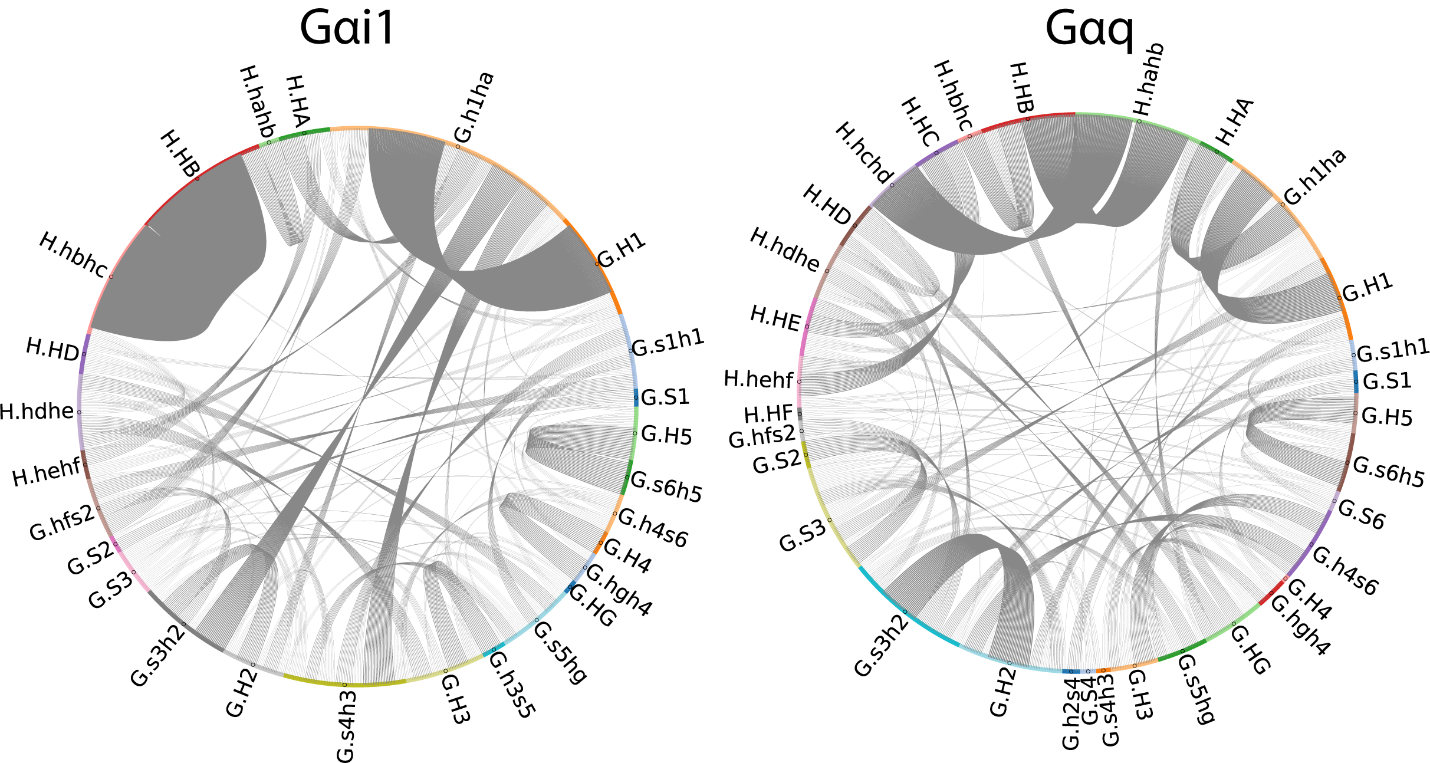

**Figure S6**. Circos map of correlated movements in Gαi1 (left) and Gαq (right). Each arch represents one structural region of Gα subunit, starting with G.S1 and going counterclockwise. The length of the arch is proportional to the number of residues there are in the region. Each line represents an edge in BNM. The thickness of the line is proportional to the edge weight of the represented edge.

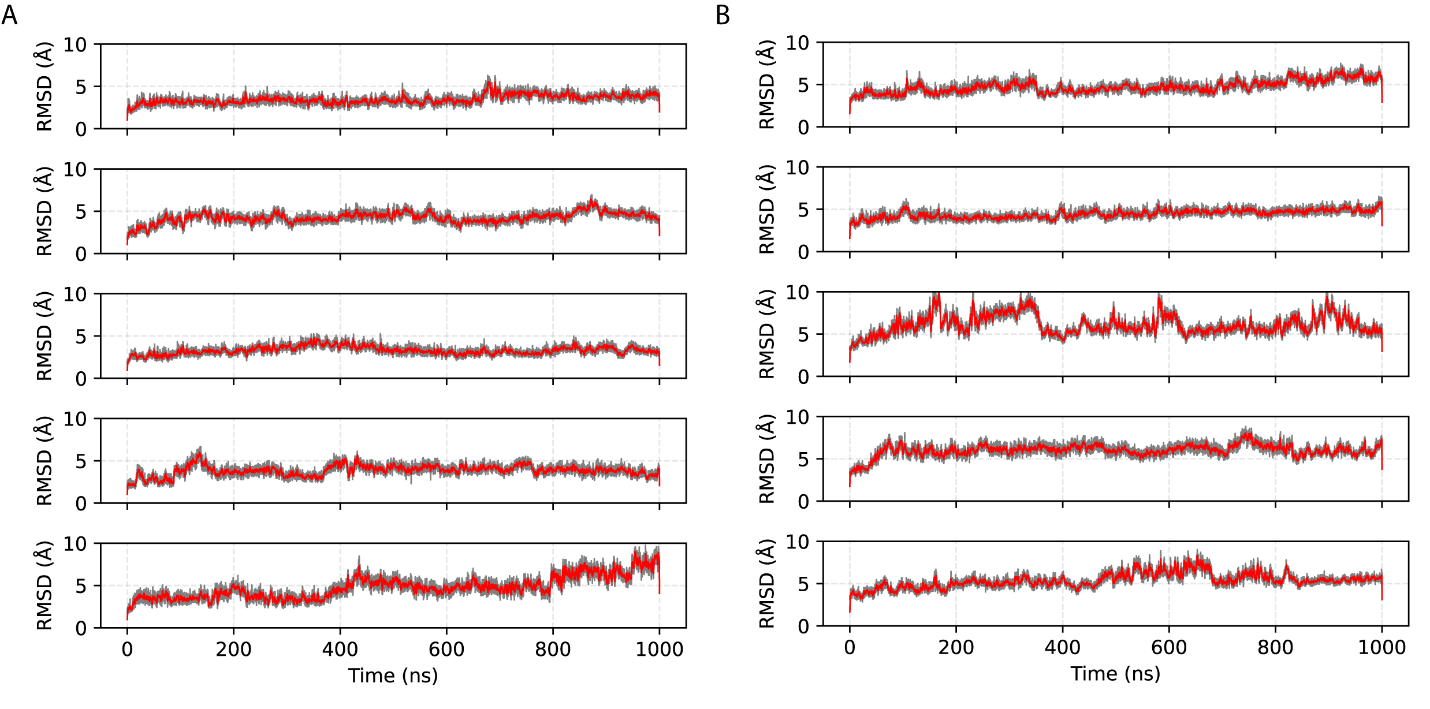

**Figure S7**. MD trajectory convergence of Gαi1 (left) and Gαq (right). Backbone atoms of Gα, Gβ, and Gγ residues were included in the calculation, The reference structure used was the pre-equilibrated structure.

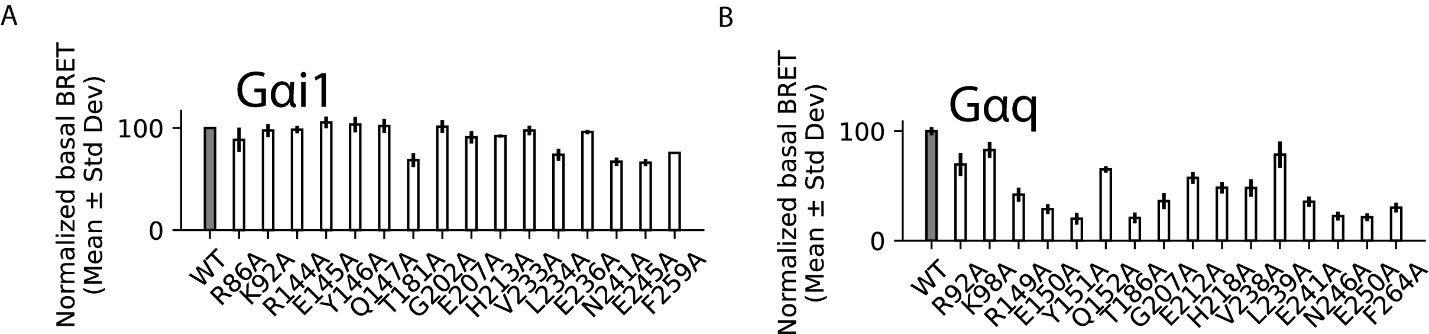

**Figure S8**. Basal BRET values for mutants, normalized to that of the wildtype.

**Table S1**. Ortholog sequence conservation of subtype specific hotspot residues.

| Common G protein Numbering | Gαi1 residue number | Gαq residue number | Gαi1 conservation percentage | Gαq conservation percentage |
| --- | --- | --- | --- | --- |
| H.HD.11 | R144 | R149 | 99.00% | 97.80% |
| H.hdhe.01 | Y146 | Y151 | 99.00% | 96.70% |
| H.hdhe.02 | Q147 | Q152 | 99.00% | 98.90% |
| G.hfs2.07 | G183 | G188 | 97.10% | 98.90% |
| G.S2.01 | I184 | I189 | 98.10% | 98.90% |
| G.s3h2.01 | G202 | G207 | 98.10% | 100.00% |
| G.H2.01 | R205 | R210 | 99.00% | 100.00% |
| G.H2.02 | S206 | S211 | 99.00% | 100.00% |
| G.H2.03 | E207 | E212 | 99.00% | 100.00% |
| G.H2.06 | K210 | K215 | 99.00% | 100.00% |
| G.H2.09 | H213 | H218 | 99.00% | 100.00% |
| G.h2s4.02 | E216 | E221 | 99.00% | 100.00% |
| G.s4h3.07 | V233 | V238 | 97.10% | 98.90% |
| G.s4h3.08 | L234 | L239 | 98.10% | 98.90% |
| G.s4h3.10 | E236 | E241 | 98.10% | 98.90% |

**Table S2**. Gαq and Gαi1 residues in its interfaces to GPCR, Ric8, adenyl cyclase, and RGS.

| Protein Partner | Interface residues |
| --- | --- |
| GPCR | G.H1.08, G.H1.11, G.H1.12, G.H2.01, G.H2.02, G.H2.03, G.H2.04, G.H2.05, G.H2.06, G.H2.07, G.H2.08, G.H2.09, G.H2.10, G.H3.01, G.H3.03, G.H3.04, G.H3.07, G.H3.08, G.H3.10, G.H3.11, G.H3.12, G.H3.14, G.H3.15, G.H3.16, G.H3.17, G.H4.04, G.H4.05, G.H4.08, G.H4.09, G.H4.12, G.H4.13, G.H4.16, G.H4.17, G.H5.02, G.H5.05, G.H5.06, G.H5.08, G.H5.09, G.H5.10, G.H5.11, G.H5.12, G.H5.13, G.H5.15, G.H5.16, G.H5.17, G.H5.18, G.H5.19, G.H5.20, G.H5.21, G.H5.22, G.H5.23, G.H5.24, G.H5.25, G.H5.26, G.HG.03, G.HG.06, G.HG.10, G.HG.12, G.HG.15, G.HG.17, G.HN.28, G.S1.01, G.S1.02, G.S1.03, G.S1.07, G.S2.01, G.S2.02, G.S2.03, G.S3.01, G.S3.02, G.S3.03, G.S3.04, G.S3.06, G.S3.08, G.S5.01, G.S6.01, G.S6.02, G.S6.03, G.S6.04, G.S6.05, G.h1ha.01, G.h1ha.02, G.h2s4.01, G.h2s4.02, G.h2s4.03, G.h2s4.04, G.h2s4.05, G.h3s5.01, G.h3s5.02, G.h4s6.02, G.h4s6.03, G.h4s6.04, G.h4s6.08, G.h4s6.09, G.h4s6.10, G.h4s6.11, G.h4s6.12, G.h4s6.13, G.h4s6.20, G.hfs2.03, G.hfs2.04, G.hfs2.05, G.hfs2.06, G.hfs2.07, G.hgh4.01, G.hgh4.02, G.hgh4.03, G.hgh4.04, G.hgh4.07, G.hgh4.08, G.hgh4.09, G.hgh4.11, G.hgh4.12, G.hgh4.13, G.hns1.01, G.hns1.02, G.hns1.03, G.s1h1.02, G.s1h1.03, G.s1h1.05, G.s1h1.06, G.s2s3.01, G.s2s3.02, G.s3h2.02, G.s3h2.03, G.s4h3.07, G.s4h3.08, G.s4h3.12, G.s4h3.13, G.s4h3.14, G.s4h3.15, G.s6h5.04 |
| Ric8 | G.H1.04, G.H1.05, G.H1.07, G.H1.08, G.H2.01, G.H2.04, G.H2.06, G.H2.07, G.H2.08, G.H2.09, G.H3.02, G.H3.04, G.H3.05, G.H3.08, G.H3.11, G.H3.12, G.H3.15, G.H3.16, G.H3.17, G.H3.18, G.H4.03, G.H4.04, G.H4.07, G.H4.08, G.H4.09, G.H4.12, G.H5.03, G.H5.04, G.H5.05, G.H5.06, G.H5.07, G.H5.08, G.H5.10, G.H5.11, G.H5.12, G.H5.13, G.H5.14, G.H5.15, G.H5.16, G.H5.18, G.H5.19, G.H5.20, G.H5.21, G.H5.22, G.H5.23, G.H5.24, G.H5.25, G.H5.26, G.HG.01, G.S1.01, G.S1.02, G.S1.04, G.S1.07, G.S2.06, G.S2.08, G.S3.01, G.S3.02, G.S3.03, G.S3.05, G.S3.07, G.S4.01, G.S4.04, G.S4.07, G.S5.01, G.S5.03, G.S5.05, G.S6.02, G.S6.03, G.S6.04, G.S6.05, G.h2s4.02, G.h2s4.05, G.h3s5.01, G.h3s5.02, G.h4s6.12, G.h4s6.20, G.hgh4.15, G.hgh4.16, G.hgh4.18, G.hgh4.20, G.hns1.03, G.s1h1.01, G.s1h1.02, G.s1h1.03, G.s2s3.01, G.s3h2.01, G.s3h2.02, G.s6h5.01, G.s6h5.02, G.s6h5.04, G.s6h5.05 |
| RGS | G.H1.09, G.H2.02, G.H2.03, G.H2.05, G.H2.06, G.H2.09, G.S2.01, G.S2.02, G.S2.04, G.hfs2.03, G.hfs2.04, G.hfs2.05, G.hfs2.06, G.hfs2.07, G.s3h2.03, G.s4h3.09, G.s4h3.10, G.s4h3.11, G.s4h3.12, H.HA.03, H.HA.06, H.HA.09, H.HA.10, H.HA.13, H.HA.24, H.HB.13, H.hbhc.01, H.hbhc.03, H.hbhc.04 |

Table S3. PDB IDs used for interface identifications.

**GPCRs**: 3SN6, 5UZ7, 5VAI, 6B3J, 6CMO, 6D9H, 6DDE, 6DDF, 6E3Y, 6G79, 6GDG, 6K41, 6K42, 6KPF, 6KPG, 6LFM, 6LFO, 6LI3, 6LMK, 6LML, 6LPB, 6M1H, 6M1I, 6N4B, 6NI3, 6NIY, 6OIK, 6OMM, 6ORV, 6OS9, 6OSA, 6OT0, 6P9X, 6P9Y, 6PT0, 6UUN, 6UUS, 6UVA, 6VCB, 6VMS, 6VN7, 6WHC, 6WI9, 6WPW, 6WWZ, 6WZG, 6X18, 6X19, 6X1A, 6XBJ, 6XBK, 6XBL, 6XBM, 7AUE, 7BB6, 7BB7, 7BW0, 7BZ2, 7C2E, 7CFM, 7CFN, 7CKW, 7CKX, 7CKY, 7CKZ, 7CMU, 7CMV, 7CRH, 7CX2, 7CX3, 7CX4, 7CZ5, 7D3S, 7D68, 7D76, 7D77, 7D7M, 7DB6, 7DFL, 7DH5, 7DHI, 7DHR, 7DTY, 7DUQ, 7DUR, 7DW9, 7E14, 7E2X, 7E2Y, 7E2Z, 7E32, 7E33, 7E9G, 7E9H, 7EB2, 7EJ0, 7EJ8, 7EJA, 7EJK, 7EJX, 7EO2, 7EO4, 7EPT, 7EUO, 7EVM, 7EVY, 7EVZ, 7EW0, 7EW1, 7EW2, 7EW3, 7EW4, 7EW7, 7EXD, 7EZH, 7F0T, 7F16, 7F1O, 7F1Q, 7F1R, 7F1S, 7F1Z, 7F23, 7F24, 7F4D, 7F4F, 7F4H, 7F4I, 7F53, 7F54, 7F55, 7F58, 7F6G, 7F6H, 7F6I, 7F8V, 7F8W, 7FIM, 7FIN, 7FIY, 7JHJ, 7JJO, 7JOZ, 7JV5, 7JVP, 7JVR, 7KI0, 7KI1, 7L0P, 7L0Q, 7L0R, 7L0S, 7LCI, 7LD3, 7LD4, 7LLL, 7LLY, 7MBX, 7MTS, 7NA7, 7NA8, 7O7F, 7PIU, 7PIV, 7RBT, 7RG9, 7RKF, 7RKM, 7RKN, 7RKX, 7RKY, 7RMH, 7RTB, 7S0F, 7S1M, 7S3I, 7S8M, 7S8O, 7SBF, 7SCG, 7SQO, 7T10, 7T11, 7T2G, 7T2H, 7T6B, 7T6S, 7T6T, 7T6U, 7T6V, 7T8X, 7T90, 7T94, 7T96, 7T9I, 7T9N, 7TD0, 7TD1, 7TD2, 7TD3, 7TD4, 7TRK, 7TRP, 7TRQ, 7TRS, 7TRY, 7TUZ, 7TYF, 7TYH, 7TYI, 7TYL, 7TYN, 7TYO, 7TYW, 7TYX, 7TYY, 7TZF, 7U2K, 7U2L, 7UTZ, 7V35, 7V68, 7V69, 7V6A, 7V9L, 7V9M, 7VBH, 7VBI, 7VDH, 7VDL, 7VDM, 7VFX, 7VGX, 7VGY, 7VGZ, 7VH0, 7VIE, 7VIF, 7VIG, 7VIH, 7VKT, 7VL8, 7VL9, 7VLA, 7VQX, 7VUG, 7VUH, 7VUI, 7VUJ, 7VUY, 7VUZ, 7VV3, 7VV5, 7VVJ, 7VVK, 7VVL, 7VVM, 7VVN, 7VVO, 7W0L, 7W0M, 7W0N, 7W0O, 7W0P, 7W2Z, 7W3Z, 7W40, 7W53, 7W55, 7W56, 7W57, 7W6P, 7W7E, 7WBJ, 7WCM, 7WCN, 7WF7, 7WIC, 7WIG, 7WJ5, 7WQ3, 7WU2, 7WU3, 7WU4, 7WU5, 7WU9, 7WUQ, 7WV9, 7WVU, 7WVV, 7WVW, 7WVX, 7WVY, 7WYB, 7WZ4, 7X2C, 7X2D, 7X2F, 7X2V, 7X5H, 7X8R, 7X8S, 7X9A, 7X9B, 7X9C, 7X9Y, 7XA3, 7XAT, 7XAU, 7XAV, 7XBD, 7XBW, 7XBX, 7XJH, 7XJI, 7XJJ, 7XJL, 7XK2, 7XK8, 7XKD, 7XKF, 7XMR, 7XMS, 7XMT, 7XOU, 7XOV, 7XOW, 7XP4, 7XP5, 7XP6, 7XT8, 7XT9, 7XTA, 7XTB, 7XTC, 7XTQ, 7XW5, 7XW6, 7XW9, 7XWO, 7XXH, 7XXI, 7XY6, 7XY7, 7XZ5, 7XZ6, 7Y12, 7Y15, 7Y1F, 7Y24, 7Y35, 7Y36, 7Y3G, 7Y64, 7Y65, 7Y66, 7Y67, 7Y89, 7YAC, 7YAE, 7YJ4, 7YK6, 7YK7, 7YKD, 7YON, 7YOO, 7YP7, 7YS6, 7YU3, 7YU5, 7YU6, 7YU7, 7YU8, 8DCR, 8DCS, 8DZP, 8DZQ, 8DZR, 8DZS, 8E3X, 8E3Y, 8E3Z, 8EF5, 8EF6, 8EFB, 8EFL, 8EFO, 8EFQ, 8F0J, 8F0K, 8F2A, 8F2B, 8F76, 8F7Q, 8F7R, 8F7S, 8F7W, 8F7X, 8FEG, 8FLQ, 8FLR, 8FLS, 8FLT, 8FLU, 8FU6, 8FX5, 8G05, 8G59, 8G94, 8GCM, 8GCP, 8GD9, 8GDA, 8GDB, 8GDC, 8GHV, 8GUQ, 8GUR, 8GUS, 8GUT, 8GW8, 8H0P, 8H0Q, 8H2G, 8H8J, 8HA0, 8HAF, 8HAO, 8HBD, 8HCQ, 8HCX, 8HIX, 8HJ0, 8HJ1, 8HJ2, 8HJ5, 8HK2, 8HK3, 8HK5, 8HMP, 8HMV, 8HN8, 8HNK, 8HNL, 8HNM, 8HOC, 8HPT, 8HQC, 8HQE, 8HQM, 8HQN, 8HS3, 8HSC, 8HTI, 8HVI, 8I95, 8I97, 8I9A, 8I9L, 8I9S, 8IA2, 8IA7, 8IA8, 8IBU, 8IBV, 8IC0, 8ID3, 8ID4, 8ID6, 8ID8, 8ID9, 8IHB, 8IHF, 8IHH, 8IHI, 8IHJ, 8IJ3, 8IJA, 8IJB, 8IJD, 8IRR, 8IRS, 8IRT, 8IRU, 8IRV, 8ITF, 8ITL, 8ITM, 8IUK, 8IUL, 8IUM, 8IW4, 8IW9, 8IY5, 8IYS, 8IZB, 8J18, 8J19, 8J1A, 8J6D, 8J6J, 8J6P, 8J6Q, 8J6R, 8JD3, 8JD5, 8JD6, 8JGB, 8JGG, 8JHY, 8JII, 8JIL, 8JIM, 8JIP, 8JIQ, 8JIR, 8JIS, 8JIT, 8JIU, 8JLO, 8JLP, 8JLZ, 8JR9, 8JSP, 8JZ7, 8JZZ, 8K2X, 8K4N, 8KH5, 8PM2, 8SAI, 8SG1, 8U1U, 8U26, 8UHB, 8W87, 8W88, 8W89, 8W8A, 8W8B, 8W8Q, 8WC3, 8WC4, 8WC5, 8WC6, 8WC7, 8WC8, 8WCA, 8WRB, 8XBH

**Ric8**: 8EL7, 8EL8, 6VU8, 6VU5, 6TYL, 6UKT

**RGS**: 5DO9, 4EKC, 4EKD, 3C7K, 2ODE, 2IHB, 2V4Z, 2IK8, 2GTP, 1AGR
